# Toward Improving Immunotolerance for Stem Cell-Derived Islets

**DOI:** 10.1101/2022.09.15.508091

**Authors:** Quan Zhou, Hongfei Li, Dario Gerace, Igor Nikolskly, Xi Wang, Jennifer Kenty-Ryu, Jingping Zhang, Matthew Hinderhofer, Elaine Robinson, Douglas A. Melton

## Abstract

Transplanting human stem cell-derived islets (SC-islets) is a promising therapy for insulin-dependent diabetes. While functional SC-islets have been produced for clinical application, immune rejection by the host remains a challenge. Present attempts, including chronic immunosuppression and/or physical encapsulation, have some disadvantages. Here we explore a strategy to induce an immune-tolerant environment based on the immune privilege observed in the male gonad. Sperm appears after the maturation of the immune system and development of systemic self-tolerance and the testis protects these autoreactive germ cells by the physical structure of blood-testis-barrier (BTB) and active local immunosuppression. Human SC-islets transplanted into the mouse testis can be physically protected by the BTB and we find that the testis secretes cytokines that induce a population of regulatory T cells (T_regs_) that express both CD4 and CD8. We identified cytokines secreted by testis and used a cocktail of IL-2, IL-10, and TGF-β for *in vitro* co-culture and *in vivo* transplantation demonstrating improved survival of SC-islets and the induction of T_regs_.

**One Sentence Summary:** Inducing local immunotolerance by suppressive cytokines for islet Transplantation.

## INTRODUCTION

The ability to generate functional pancreatic endocrine cells from stem cells presents new options for cell replacement therapy for people with diabetes. *In vitro* differentiation of insulin-producing beta cells and glucagon-producing alpha cells alongside other cell types has been well characterized(*1–5*). While there is room to improve the final cell composition of the SC-islets, a major challenge for cell therapy becomes rejection of foreign cells by the recipient’s immune system. Projects aimed at preventing allo-rejection include human leukocyte antigen (HLA) haplotype bank or hypoimmunogenic human induced pluripotent stem cells (IPSCs)(*6–9*). Here we present evidence for a complementary approach by exploring naturally existing immune privilege in the gonad and reconstructing a tolerant microenvironment for prolonged survival of the human SC-islet grafts.

The mammalian testis is a well-studied site of immune privilege(*10*). The onset of spermatogenesis starts after puberty, representing a unique challenge to the immune system as neoantigens from meiotic and haploid germ cells appear after the formation of systemic self-tolerance. To protect immunogenic germ cells from systemic immune attack, testicular immune privilege is maintained by coordinating systemic immune tolerance, the local physical structure, and active local immunosuppression(*11*). The physical structure, the blood-testis barrier (BTB), divides the seminiferous epithelium into the basal and the abluminal compartments. This separates the host immune cells from meiosis and spermiogenesis which take place in a specialized microenvironment behind the BTB(*12*). In addition, Sertoli cells and Leydig cells, which secrete immunosuppressive factors, can directly or indirectly suppress immune cell activation. Consequently, the testis is a privileged environment wherein both allo- and auto-antigens can be tolerated without immune rejection(*13–15*).

Here we explore the protection of human stem cell-derived islets (SC-islets) by taking clues from the testis microenvironment. SC-islets were transplanted into the seminiferous tubules (inside the BTB) or interstitium (outside the BTB) of mouse testes. SC-islets had up to eight weeks of survival both inside and outside the mouse BTB. However, beta cells only accurately regulated blood sugar when transplanted into the interstitium of the testis. In an *in vitro* co-culture assay of the human SC-islets with C57BL/6 mouse mononuclear cells (MNCs), we observed suppressed T cell activation and proliferation in the presence of testis tissue or Sertoli cells. We identified a specific group of T_regs_ with both CD4 and CD8 expression and showed that they can protect human SC-islet in *vitro* co-culture with mouse MNCs and in vivo transplantation. We further show that IL-2, IL-10, and TGFβ secreted by testis cells can induce T_regs_ and suppress T cell activity. The cytokine cocktail induces an immunotolerant local environment by activating T_regs_ and may provide a path forward to protect SC-islets from immune rejection.

## RESULTS

### Both Inside and Outside the BTB Are Protective Environment

Human embryonic stem cells (hESCs) were differentiated into SC-islets through a stepwise protocol^1^. Within SC-islets, which contain several endocrine cell types, beta cells were characterized by NKX6-1 and C-peptide expression at day 14 in the last stage of differentiation (Fig. S1A). SC-islets were resized to 100mm in diameter (Fig. S1B) and their function was assessed by glucose-stimulation-insulin secretion (GSIS) (Fig. S1C). SC-islets were transplanted into C57BL/6 male mice testis, both inside and outside the seminiferous tubules (Fig. S1D). Note the mice have functional beta cells that can regulate their blood glucose uptake. We measured human insulin levels to evaluate SC-islet existence and function. Following transplantation of SC-islets, human insulin levels were assessed in mouse serum and by in vivo GSIS. Human insulin levels began to rise two weeks after transplantation and remained steady for about six weeks both inside and outside the seminiferous tubules (Fig. 1A). The human insulin levels observed for SC-islets transplanted in the interstitium were higher than in the seminiferous tubules, perhaps due to the tight junction surrounding the tubules which may slow or block the insulin release into the serum. In vivo GSIS of the recipient mice at four weeks post transplantation showed that SC-islets are functional when transplanted into the testis interstitium; less so when transplanted into seminiferous tubules of the testis (Fig. 1B).

**Figure1.**
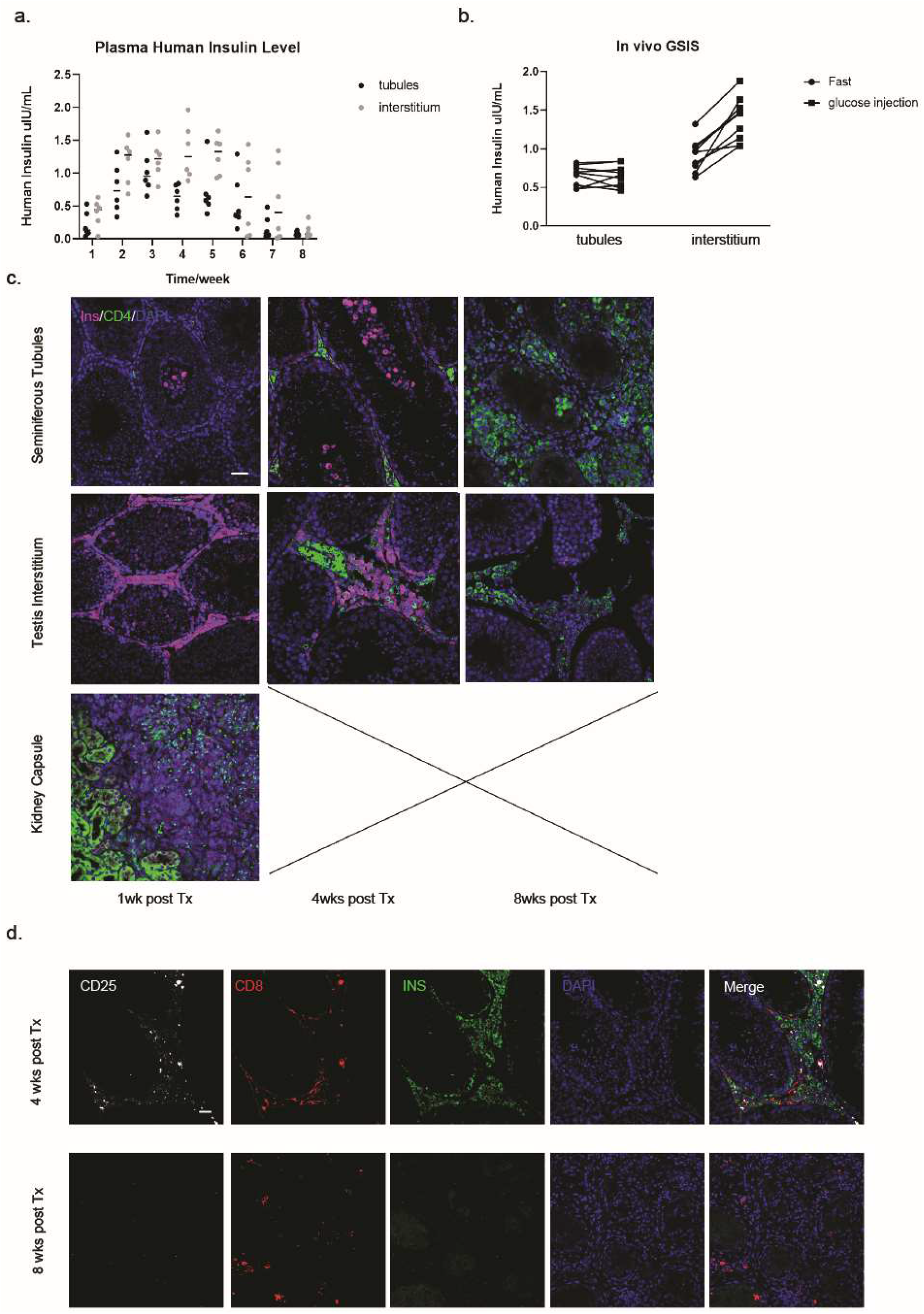
Human SC-islets survival and function in C57BL/6 mice testes A. Human insulin level in mouse plasma measured every week after transplantation. B. Glucose-stimulation-insulin-secretion *in vivo*, measured four weeks after transplantation. C. Immunostaining of INS and CD4 in the testes (or kidney capsules) at 1, 4, and 8 weeks after transplantation. No grafts were found 1 week after transplantation under the kidney capsule (bar =100μm). D. Immunostaining of CD25, CD8, and INS, showing T_regs_ in the testes 4 and 8 weeks after transplantation (bar =100μm).

Survival and immune cell infiltration in grafts were assessed by immunostaining of SC-β cells and mouse T cells. Transplanted SC-islets survived at least four weeks in both the seminiferous tubules and the interstitium, whereas SC-islets are eliminated within one week after transplantation to the kidney capsule (Fig 1C). Infiltration by mouse CD4^+^ T cell is evident in the kidney capsule in the first week after transplantation, but not in the testis (both in and out the seminiferous tubules) until four weeks post-transplantation. SC-islets secreted insulin four weeks after transplantation in the testis but by eight weeks they were rejected in both interstitium and seminiferous tubules. In the seminiferous tubules transplantation group, the CD4^+^ T cells were found inside and eliminated both SC-islets and all the mouse germ cells, showing that the germ cells are autoreactive to the immune system and the physical barrier is crucial to protect them from immune rejection. In contrast, when SC-islets were transplanted into the interstitium of the testis, the T cells remained outside the seminiferous tubules and only killed transplanted grafts. Staining for CD25 and CD8 (T_regs_) at four weeks after transplantation, shows T_regs_ in the interstitium of the testis (Fig. 1D) but not by eight weeks. The T_regs_ in the testis are associated with prolonged survival of SC-islets indicating a possible role in immune tolerance in testis.

### CD4^+^/CD8^+^ Cells Provide Immune Suppression *in vitro*

To further assess the immune suppression observed in testis, MNCs from mouse spleen were cultured with SC-islets in the absence or presence of testis. We set up the following six conditions (Fig. 2A): 1. A control of mouse MNCs alone; 2. MNCs with SC-islets; 3. MNCs with seminiferous tubules tissues; 4. SC-islets injected into testis then removed and co-cultured with MNCs, to mimic *in vivo* transplantation into the seminiferous tubules of the testis; 5. SC-islets injected outside of the seminiferous tubules and co-cultured with MNCs, to mimic the *in vivo* transplantation into the interstitium spaces of the testis, 6. SC-islets co-cultured with MNCs in the presence of Sertoli cells. CD4^+^ T cell activation was measured by CD62L expression in the CD3^+^/CD4^+^ T cell population and down-regulation of CD62L indicates T cell activation. The results show that more than 90% of the CD3^+^/CD4^+^ T cells remain inactivated in the control of MNCs only, whereas more than 50% of the CD3^+^/CD4^+^ T cells are activated and stimulated by human SC-islets co-cultured with MNCs. The other four test groups, which contained either seminiferous tubules or Sertoli cells, show suppressed T cell activation (Fig. 2B). T cell proliferation was measured three days after co-culture and was suppressed in the presence of testis tissue or Sertoli cells (Fig. 2B). These results indicate that activation and proliferation of CD3^+^/CD4^+^ T cells is suppressed by the testis, likely by Sertoli cells, and not only by the physical barriers of BTB.

**Figure2.**
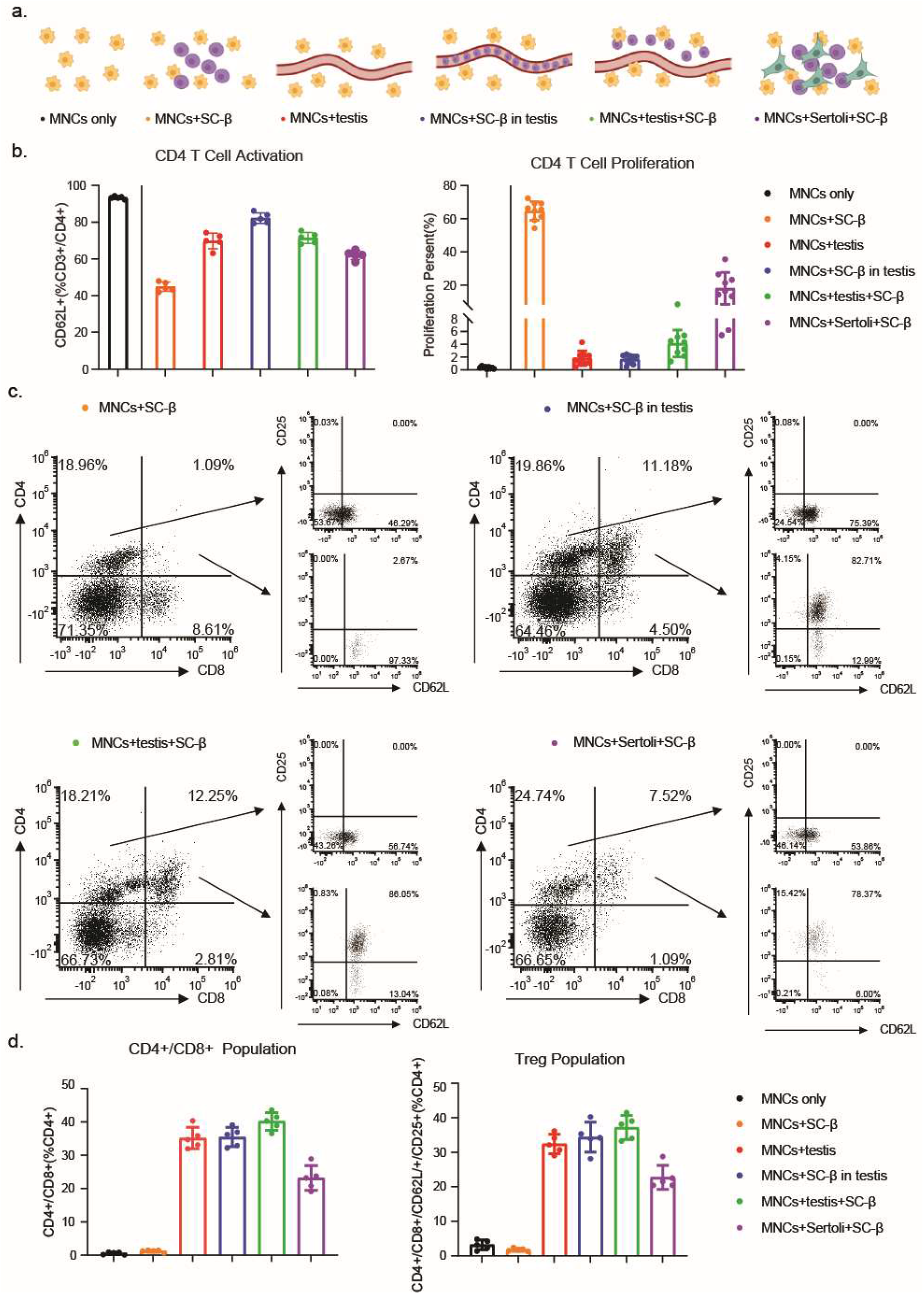
*In vitro* co-culture of SC-islets and MNCs in the presence of Testis tissue or Sertoli cells A. Schematic diagram of co-culture settings. B. CD4 T cell activation (left) and proliferation (right) under co-culture. C. Flow cytometry shows CD4^+^/CD8^+^ cells exist in co-culture groups containing testis tissues or Sertoli cells. This population has high CD25 and CD62L expressions. D. Statistical analysis of the ratio of CD4^+^/CD8^+^ and CD25^+^ population in each co-culture condition.

We measured the activity of CD8^+^ cytotoxic T cells by co-staining for CD8 and CD4 in the CD3 positive population. We find a unique T cell population with the presence of testis tissue or Sertoli cells (Fig. 2C). This CD4^+^/CD8^+^ population is CD25 positive, consistent with a T_re_g identity. Nearly 50% of the CD4^+^ T cells are CD4^+^/CD8^+^ and 90% expressed CD25 (Fig. 2D). In conclusion, testis tissue or Sertoli cells induce a CD4^+^/CD8^+^ T cell population in *vitro*.

### CD4^+^/CD8^+^ Population Exists *in vivo* and Plays an Immunosuppressive Role

Then we asked whether the CD4^+^/CD8^+^ population exists *in vivo* and might have a role in inducing tolerance. CD4^+^/CD8^+^ cells are observed four weeks after human SC-islets transplantation into the testis (Fig. 3A, 3B). The CD4^+^/CD8^+^ cells only appear in the interstitium of the testis, whereas CD8 single positive cytotoxic cells migrate into the seminiferous tubules (Fig. 3A). CD4^+^/CD8^+^ cells and CD4 single-positive cells were separated by flow cytometry (Fig. 3B) and express mRNA encoding FOXP3 as well as the suppressive cytokines TGFβ and IL-10. CD4 single-positive cells show a different cytokine profile including IL-2, IL-4, IL-17, INFγ, and TNFα (Fig. 3C). These results are consistent with CD4^+^/CD8^+^ cells serving an immunosuppressive role (Fig. 3D).

**Figure 3.**
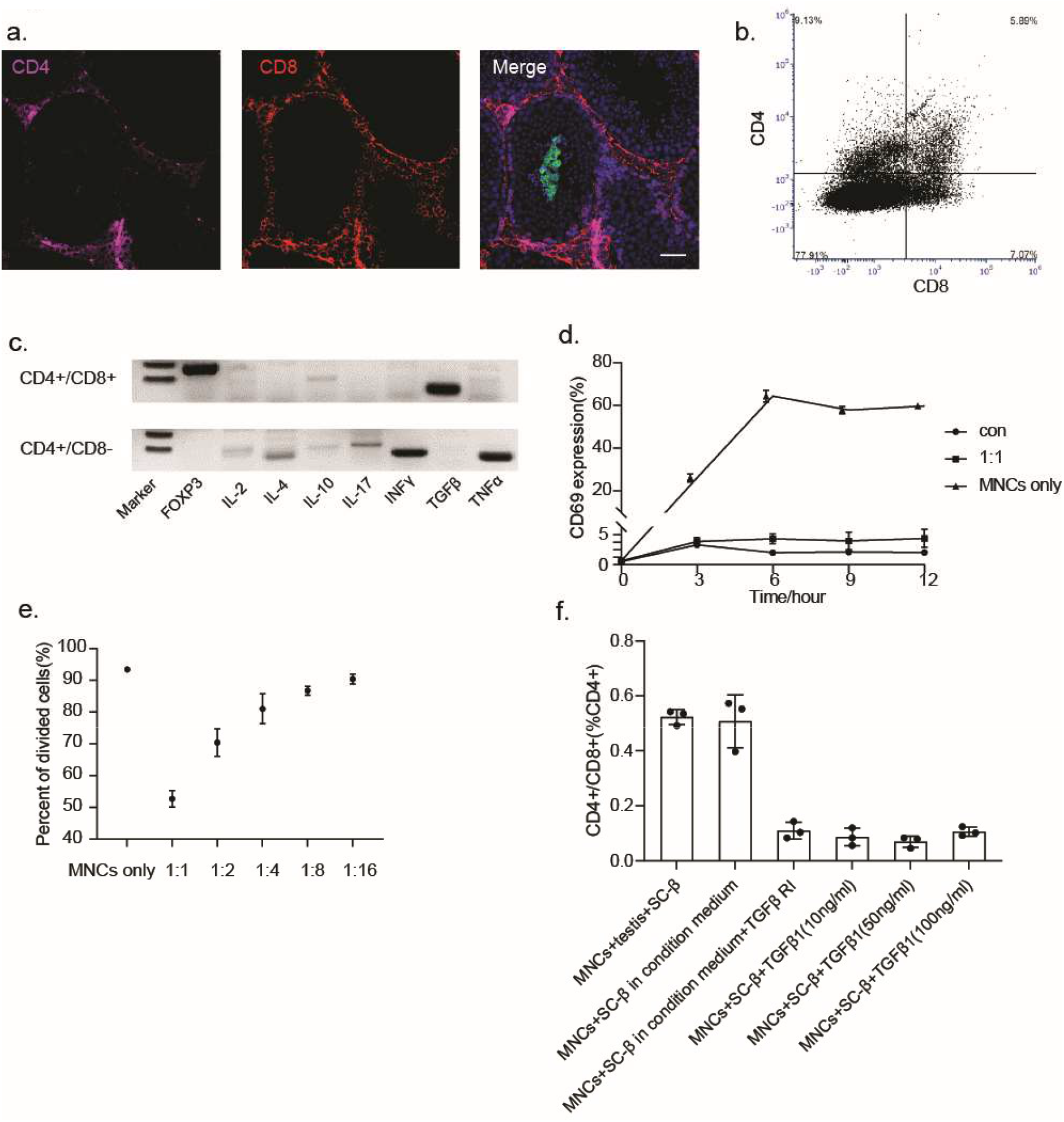
Identification of CD4^+^/CD8^+^ population in testis transplantation A. Immunostaining showing CD4^+^/CD8^+^ cells in the interstitium of testis 2 weeks after transplantation. B. FACS sorting of CD4^+^/CD8^+^ cells from transplanted testis 2 weeks after transplantation. C. RT-PCR shows cytokines produced by CD4^+^/CD8^+^ and CD4^+^/CD8^-^ populations 4 weeks after transplantation. D. T cells activation stimulated by CD3/CD28 antibody in the presence of CD4^+^/CD8^+^ cells isolated from transplanted testis 2 weeks after transplantation. The control group was MNCs without CD3/CD28 antibody or CD4^+^/CD8^+^ cells. The MNC only group stimulated by CD3/CD28 antibodies. The 1:1 group contains MNCs and CD4^+^/CD8^+^ cells at a 1:1 ratio, then stimulated by CD3/CD28 antibodies. E. T cell proliferation in CD3/CD28 antibody stimulation assay in presence of CD4^+^/CD8^+^ cells at CD4^+^/CD8^+^ cells: MNCs = 1:1, 1:2, 1:4, 1:8, 1:16 ratios. F. CD4^+^/CD8^+^ population induction in vitro co-culture assays of SC-islets and MNCs.

We performed bulk RNAseq of the T_regs_, CD4^+^/CD8^+^ cells, and CD4 single-positive cells isolated from SC-β transplanted testis and find that the CD4^+^/CD8^+^ cells have a gene expression profile most similar to that of T_regs_ (Fig S2A). The methylation of genes related to T_regs_ cells was assessed in the CD4^+^/CD8^+^ population (Fig. S2B) and compared to the T_regs_ isolated from mouse spleen and CD4 single-positive cells isolated from SC-β transplanted testis. CD4^+^/CD8^+^cells show significant levels of demethylation of CTLA4, FOXP3, IKZF2, IKZF4, IL-2Rα, and TNFRSF18, matching that of T_reg_ cells. The methylation levels of PDL1, CDKN1C, IL-2, IL-4, and TNF were similar among all the samples. The methylation levels of these genes in the CD4^+^/CD8^+^ population were measured at different time points (two weeks, four weeks, and eight weeks) after transplantation revealing a gradual change in the methylation of CTLA4, FOXP3, IKZF2, IKZF4, IL-2Rα, and TNFRSF18, indicating that the CD4^+^/CD8^+^ cells lose their identity as T_regs_ by the time of graft rejection.

To test whether the CD4^+^/CD8^+^ induce tolerance, CD4^+^/CD8^+^ cells were isolated from SC-islets transplanted testis 2 weeks after transplantation. Their ability to suppress T cell activation was tested using an in *vitro* T cell activation assay stimulated by CD3/CD28 beads (Fig. 3D). The results showed that co-culturing with CD4^+^/CD8^+^ cells down-regulates CD69 expression (a T cell activation marker) in a dosedependent manner. CD4^+^/CD8^+^ cells could also suppress T cell proliferation in a dose-dependent manner (Fig. 3E).

Conditioned media of testis tissue, in the MNC vs SC-islet co-culture assay, was used to test whether tolerance is provided by soluble signals. An equal ratio of the CD4^+^/CD8^+^ population is induced by the presence of testis tissue or testis condition media (Fig. 3F). TGFβ receptor inhibitor (SB-431542) was added to the testis condition media, and this reduced the CD4^+^/CD8^+^ population, but the addition of TGFβ1 to the co-cultures without testis tissue or testis condition media does not increase the CD4^+^/CD8^+^ population (Fig. 3F). These data suggest that the CD4^+^/CD8^+^ population is induced by soluble factors secreted by testis tissue and that TGFβ plays a role.

### CD4^+^/CD8^+^ Cells Help Induce Immune Tolerance When Transplanted Together with SC-islets

SC-islets were transplanted into C57BL/6 mice testis and four weeks later retrieved the transplants isolated CD4^+^/CD8^+^ cells and CD4^+^/CD8^-^ cells (Fig 4A). These two separate cell populations were reaggregated with SC-islets and transplanted into the kidney capsule in C57BL/6 mice. The SC-islets reaggregated with CD4^+^/CD8^-^ cells died one week after transplantation (Fig. 4B) whereas SC-islets reaggregated with CD4^+^/CD8^+^ cells survived for eight weeks. The CD4^+^/CD8^+^ cells remain in the grafts four weeks after transplantation as does a CD25^+^ population of cells (Fig. 4C). These results indicate that the CD4^+^/CD8^+^ cells can play a protective role in vivo when co-transplanted with SC-islets outside testis tissue.

**Figure 4.**
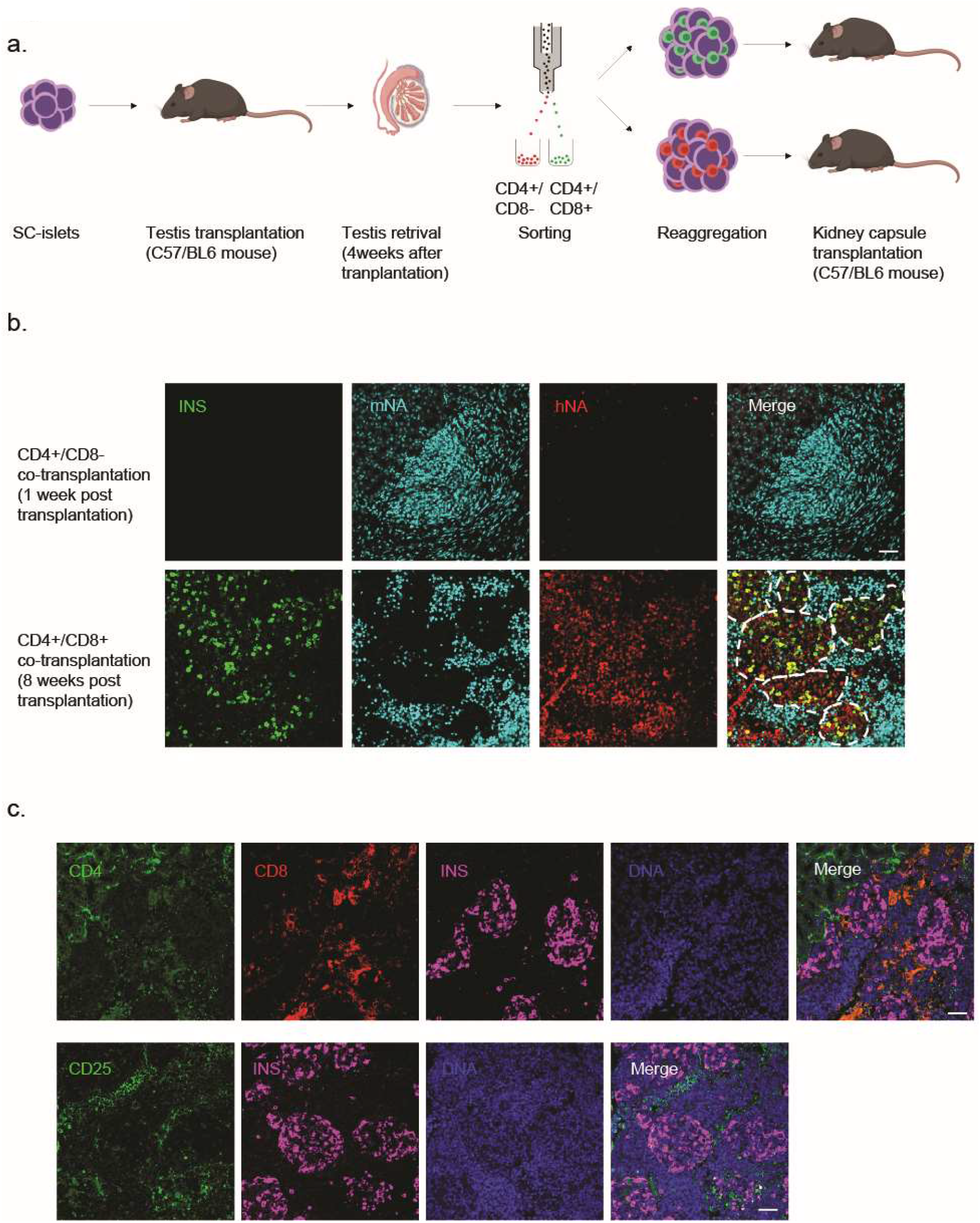
CD4^+^/CD8^+^ cells are protective for *in vivo* transplanted grafts in sites outside of the testis A. Schematic diagram showing two rounds of transplantation experiments. B. Immunostaining of insulin-producing cells, mouse resource cells, and human resource cells of CD4^+^/CD8^-^ cells co-transplantation and CD4+/CD8+ cells co-transplantation at 1 or 8 weeks after transplantation (bar = 100μm). C. Immunostaining shows CD4^+^/CD8^+^ cells and CD25^+^ T_regs_ exist 4 weeks after transplantation (bar = 100μm).

### Immunotolerant Microenvironment Induced by Cytokines Protects SC-islets both *In vitro* and *In Vivo*

We next tested for cytokines and chemokines secreted by mouse testis and identified IL-2, IL-10, CXCL1, CCL2, TIMP-1, and CXCL12 (Fig. 5A and B). Secretion of IL-2 and IL-10 in the testis condition media was confirmed by ELISA (Fig. 5C). Addition of IL-2 (30 IU/ml) induced the highest Treg proportion in the co-culture among conditions tested (Fig. 5D). IL-2 expands not only T_regs_ but also effector T cells (Teffs), natural killer cells (NKs), and cytotoxic T lymphocytes (CTLs)(*16, 17*), and IL-2 alone also up-regulated T cell proliferation in the co-cultures (Fig. 5E). We observed down-regulated T cell expansion in the activation condition with IL-10 or TGFβ (Fig. 5E). Previous literature also indicated that IL-2 and TGFβ are crucial for inducing Treg fate when T cells are stimulated(*18*), while IL-10 was an essential mediator of suppression used by T_regs_(*19, 20*). Thus, we chose IL-2, IL-10, and TGFβ as the cytokine cocktail to test further.

**Figure 5.**
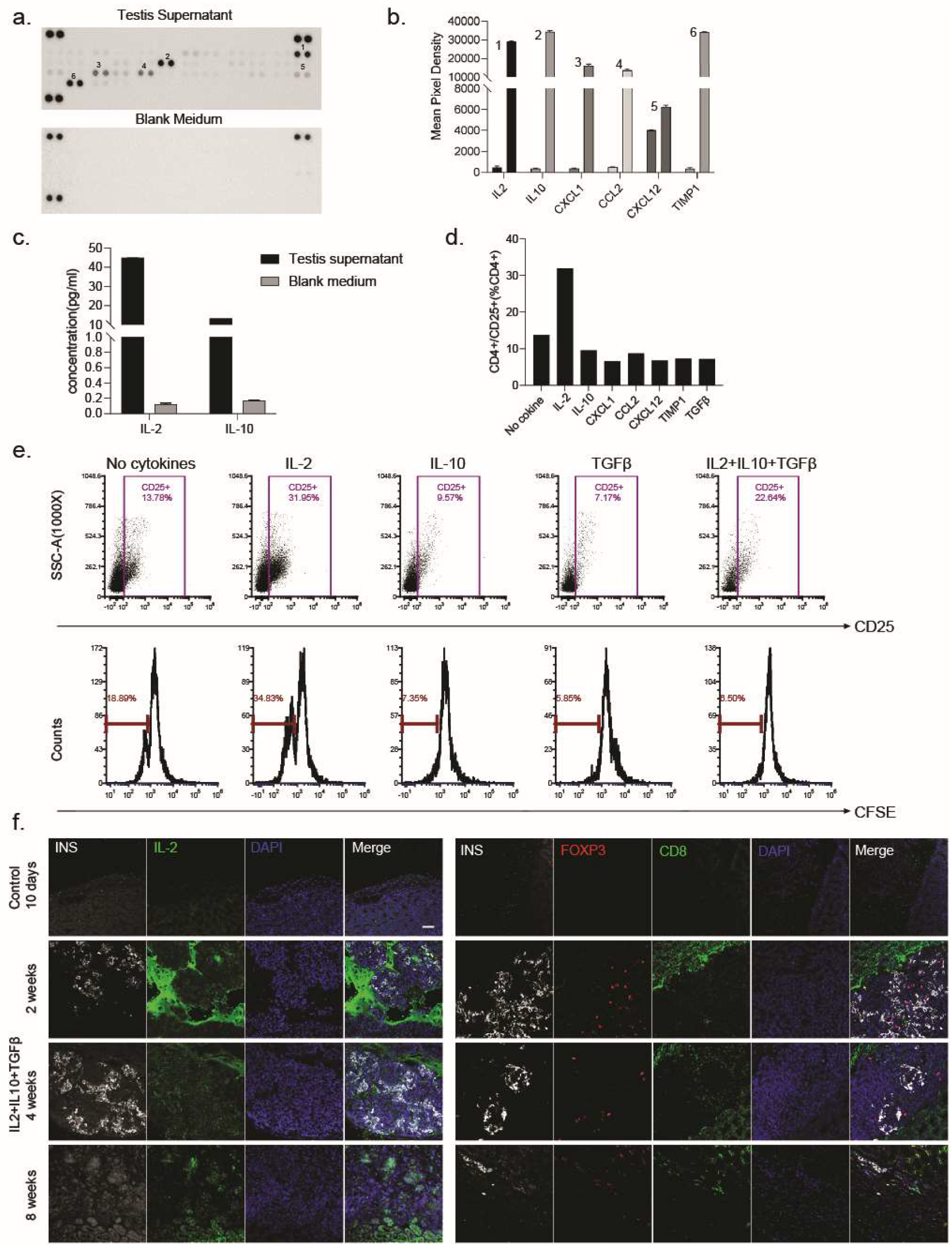
Reconstitution of the immune tolerant environment using cytokine cocktails. A. Cytokine array of secreted cytokines and chemokines in testis supernatant. The spots on the top left, top right and bottom left were the reference spots. B. Quantification of cytokines and chemokines normalized to the blank medium control. C. IL-2 and IL-10 Elisa of testis supernatant confirm secretion of cytokines in the testis. D. Cytokines screening for induction of T_regs_ in by CD3/CD28 beads. 3 days after expansion, MNCs were collected for CD25 and CD4 staining. E. T cell stimulated by CD3/CD28 beads while adding IL-2, IL-10, TGFβ, or a combination of cytokines, measured 3 days after co-culture. Upper row, T_regs_ induction under each condition. Lower row, T cell proliferation tested by CFSE cell proliferation trace under each condition. F. Immunostaining of IL-2, INS, FOXP3, and CD8 in grafts at 10 days in the control group, and 2, 4, and 8 weeks show a protective effect of the cytokine cocktails (bar = 100 μm).

T_regs_ induction and down-regulated T cell proliferation were achieved in the co-culture assay with IL-2, IL-10, and TGFβ together (Fig. 5E). In the transplantation experiments, SC-islets clusters were embedded with these cytokines in Matrigel, using control of SC-islets only. Transplants were retrieved at different times and we found that grafts without cytokines were rejected within ten days post transplantation. SC-islets embedded with cytokines had a longer life span (up to 8 weeks) under the kidney capsule in these immune-competent mice (Fig.5F).

Two weeks after transplantation there was a robust IL-2 signal in the gap space in the transplanted grafts under the kidney capsule. FOXP3^+^ T_regs_ infiltrated the grafts. Four weeks post transplantation the signal of IL-2 was diminished and fewer T_regs_ were observed in the grafts. At eight weeks no IL-2 signal nor T_regs_, were detected and the SC-islets were rejected. We could conclude that IL-2, IL-10, and TGFβ as a cytokine cocktail can form a local immunotolerant environment to induce T_regs_ and protect transplanted grafts against xeno-rejection *in vivo*.

### The cytokine cocktail protects NOD islets from auto-rejection

The experiments described above address immune protection in a xeno-tranplant. To test whether these cytokines provide protection in an auto-immune diabetic model, we injected pancreas homing AAV8 virus-derived with a mouse INS2 reporter driving a mouse IL-2 mutein (N88D) and TGFβ into three-week-old female NOD mice. The modification of IL-2 into IL-2N88D follows reports(*21, 22*) showing that reduced binding to the intermediate affinity IL-2Rβγ receptor favors Treg-specific expansion. As the AAV8 had a length limitation, IL-10 was abandoned since it did not induce T_regs_. Firefly luciferase (Luc2) was used to trace the virus location in vivo. Control mice were injected only with INS2-derived Luc2. The luciferase signal and blood glucose of all mice were measured every week (Fig. 6B and C). Blood glucose levels show the mice became diabetic at 16-18 weeks after injection in both the untreated control and luc2 control mice. In contrast, half of the mice injected with IL-2N88D+TGFβ remained normal glycemic at 20 weeks after injection. The survival rate of the mice was also higher in the IL-2N88D+TGFβ injection group (Fig. 6C). Histology of the pancreases of NOD mice from 10 weeks and 20 weeks after injection showed the absence of healthy islets in the controls (Fig. 6D). The islets were either heavily infiltrated with CD4^+^ T cells or totally rejected. In contrast, islets in the mice injected with IL-2N88D+TGFβ virus were either free of T cells or infiltrated with T_regs_. There was a lower T cell density where T_regs_ were located (Fig. 6D), indicating that T_regs_ were suppressed T cell proliferation. 50 islets were examined by immunohistochemistry in this cohort: 14 lacked any T cell infiltration and among the others, 33/36 had T_regs_. From these results, we could conclude that cytokine cocktail can recruit T_regs_ to protect islets from being rejected for some period.

**Figure 6.**
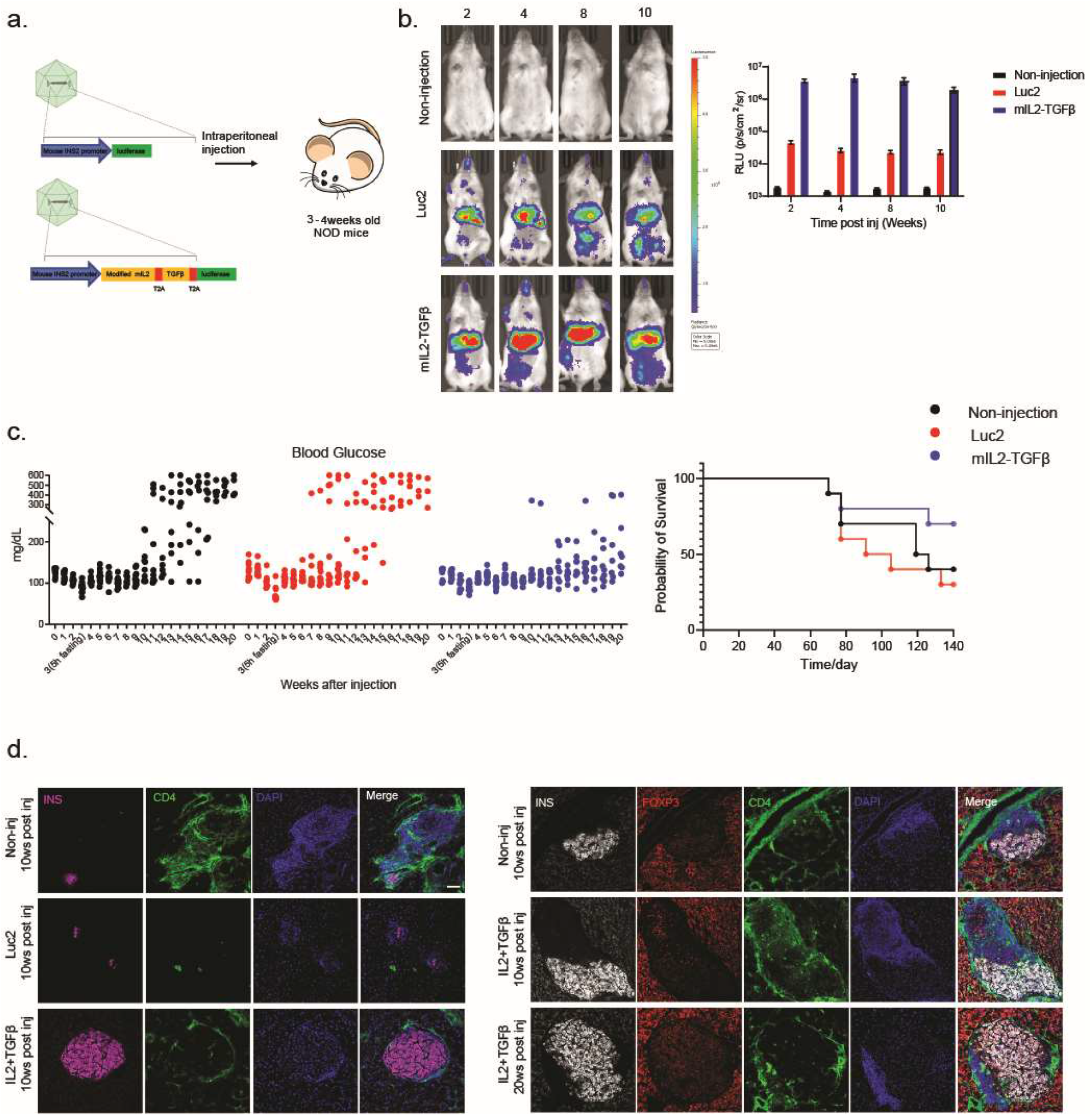
A cytokine cocktail induces a tolerant environment to help prevent or delay diabetes in NOD mice. A. Schematic diagram showing pancreas homing AAV8 derived mouse Ins2 promoter driving mIL-2N88D and TGFβ (the control group has mouse Ins2 derived Luc2 in the virus) in 3–4-week-old NOD mice. B. Luciferase signal showing virus expression *in vivo*, measured at 2-, 4-, 8- and 10-weeks post-injection. C. Blood glucose level and survival rate of each group monitored every week. The 3-week blood glucose was measured after 5 hours of fasting; the other time points were measured without fasting. D. Immunostaining shows islet survival and Treg induction.

## DISCUSSION

Stem cell-derived cells or tissues hold promise in providing an endless cell source for tissue or organ replacement. After producing fully functional stem cell-derived cells, the next challenge is solving the problem of immune rejection. In this study, we mimic the local immunotolerant environment of an immune-privileged organ to protect cells. We find a tolerizing T cell population, CD4^+^/CD8^+^ cells, in mouse testis, which has been previously described in the literature in several pathological conditions(*23–28*). In most cases, including cancer, autoimmune, and chronic inflammatory disorders, it serves a suppressive role in regulating immune activity(*26–28*). In our study, the CD4^+^/CD8^+^ cells represented T_regs_ identity in many aspects, including TSDR, gene expression, and cytokine secretion. We also find that this population of cells can suppress T cell activities and protect transplanted SC-islet grafts both in vitro and in vivo. To extend our study, we use a cytokine cocktail identified in the testis secretome (IL-2, IL-10, and TGFβ) to induce a protective environment for both xeno-transplantation and autoimmune rejection in NOD mice. The results suggest that the cytokine cocktail could induce T_regs_ population when the antigen-presenting cells are transplanted. Human pluripotent stem cell-derived islets (transplanted SC-islets) and endogenous NOD islets both survive longer in vivo. These data indicate that the cytokine cocktail can induce a protective micro-environment that may provide a possibility for improving cell allo-cell transplantation outcomes.

We show that the SC-islets could survive up to 2 months after transplantation into an immune-competent mouse. The Matrigel and the associated cytokines may dissipate when the Matrigel dissolves. A longer-lived matrix or a slow-release pump may achieve a longer life span for transplanted grafts. Generating an IL-2, IL-10, and TGFβ secreting pluripotent cell line may also be explored in which case would produce and release the tolerizing cytokines for a longer period. In order to test this hypothesis, we generated an IL-2, IL-10, and TGFβ constant expression cell line (2B10) in a separate project and used it to test the protective efficacy after SC-islet transplantation(*29*). In two mouse models (C57BL/6 and NOD), we show that the 2B10-derived SC-islets can recruit T_regs_ to their surrounding area and have a significantly longer life span compared to the WT SC-islets. Those studies extend and complement the present report wherein are directly provided to the surrounding micro-environment. Altogether all these results suggest multiple ways to deliver cytokines to provide a protective function.

Numerous studies have shown that T_regs_ play a central role in the induction and maintenance of immune homeostasis and self-tolerance. Phase I/II clinical trials have explored administering low-dose IL-2 subcutaneously to induce T_regs_ for treating autoimmune diseases such as chronic refractory graft-versus-host-disease (GVHD)(*30*), hepatitis C virus-induced vasculitis(*31*), and type 1 diabetes (T1D)(*32–34*). These studies showed that a low dose of IL-2 would yield a higher T_regs_ ratio in the peripheral blood of the patients. However, typical side effects such as influenza-like syndrome were reported(*32*), which might be due to the systematic suppression of the immunity in the patients. Here, we aim to induce the T_regs_ surrounding the grafts instead of a systematic induction of T_regs_.

Overall, our approach provides another approach to considering immune protection of stem cell-derived tissue transplantation with the aim of avoiding the long-term usage of systemic immunosuppressive drugs.

## MATERIALS AND METHODS

### Stem cell culture and islet differentiation

HUES8 human embryonic stem cells are cultured and differentiated into SC-islets in suspension as previously described(*1*). Briefly, HUES8 was maintained using mTeSR1 (Stem Cell Technologies, 85850) in 500ml spinner flasks (Corning, VWR) spinning at 70 rpm in the incubator at 37 °C, 5% CO2, and 100% humidity, and was passaged every 3 days. For differentiation, 150 million cells were seeded using 300 ml mTeSR1+10 μM Rock inhibitor (Y27632) 3 days prior to changing to differentiation media. The stepwise SC-islets differentiation was performed as the previous described(*1*), with slight modification, the brief protocol is listed below:

Stage 1 day 1: S1 medium supplemented with Activin A (100ng/ml) and CHIR99021 (14μg/ml). Stage 1 day 2-3: S1 medium supplemented with Activin A (100ng/ml).
Stage 2 day 1-3: S2 medium supplemented with KGF (50ng/ml).
Stage 3 day1-2: S3 medium supplemented with KGF (50ng/ml), LDN193189 (200nM), Sant1 (0.25μM), retinoic acid (2μM), PDBU (500nM) and Y-27632 (10μM).
Stage 4 day 1-5: S3 medium supplemented with KGF (50ng/ml), Sant1 (0.25μM) and retinoic acid (0.1μM), Y27632 (10μM) and Activin A (5ng/ml).
Stage 5 day1-3: BE5 medium supplemented with Sant1 (0.25μM), Beta Cellulin (20ng/ml), XXI (1μM), Alk5iI (10μM) and T3 (1μM). Stage 5 day5-7: BE5 medium supplemented with Beta Cellulin (20ng/ml), XXI (1μM), Alk5iI (10μM) and T3 (1μM).
Stage 6: CMRLS medium supplemented with Alk5iI (10μM) and T3 (1μM).

All experiments involving human cells were approved by the Harvard University IRB and ESCRO committees.

### Resizing, Reaggregation, and transplantation of SC-islets into mouse testis

The SC-islet clusters were resized to 100μm in diameter to be easy to transplant. Before and after resizing, the clusters were stained with NKX6-1, C-peptide, and Chromogranin A to confirm the critical gene expression of the SC-islets (Fig. S1B). The reaggregation procedure is performed to remove non-endocrine cells before transplantation as described previously(*3*). SC-islets were dissociated into single cells at Stage 6 day 14 using TryplE and seeded back into 3D suspension in Stage 6 media in 1 million/ml concentration, and then cultured in the incubator at 37 °C, 5% CO_2_ with 70 rpm agitation. After 24h, the endocrine cells would self-aggregate into clusters, whereas progenitor cells remain in the supernatant. Media changes were performed every 2 days. The reaggregated clusters were ready to transplant after 3 days. Transplantation of reaggregated clusters into testis was carried out as previously described(*35*). 2 million reaggregated clusters were injected into 8-10 weeks old C57BL/6 mice (Charles River) testis(*36*), both inside and outside of the seminiferous tubules. At the specified time after transplantation, testis containing SC-islets were dissected and fixed in 4% PFA overnight at 4 °C. The fixed testes were embedded in paraffin and sectioned for immunofluorescence staining, which was performed as described below. All animal studies were approved by the Harvard University IACUC.

### *In vivo* and *in vitro* Glucose-stimulated insulin secretion assays

Detection of human insulin level in mouse serum following a glucose challenge in SC-islets transplanted animals was conducted as described(*1*): animals were fasted for 16h overnight, and the Serum samples were collected through mandibular bleeding using a lancet (Feather; 2017-01) for detecting fast insulin levels. Then, the glucose challenge was done by intraperitoneal injection of D-(+)-glucose (2 g/1 kg body weight). Serum was collected 30min after injection. Serum was separated out using Microvettes (Sarstedt; 16.443.100) and stored at −80°C until ELISA analysis.

Cell clusters sampled from Stage 6 day 14 SC-islets were divided into four parts for GSIS assays. Krebs buffer (128 mM NaCl, 5 mM KCl, 2.7 mM CaCl2, 1.2 mM MgSO4, 1 mM Na2HPO4, 1.2 mM KH2PO4, 5 mM NaHCO3, 10 mM HEPES (Life Technologies; 15630080), 0.1% BSA in deionized water) containing 2.8mM glucose was used to wash clusters. Afterward, the clusters were loaded into 24-well plate inserts (Millicell Cell Culture Insert; PIXP01250), followed by fasting in Krebs buffer containing 2.8mM glucose for 1h. Subsequently, clusters were washed with Krebs buffer containing 2.8mM glucose and incubated with Krebs buffer containing 2.8mM glucose at 37°C for 1h, supernatant was collected to measure insulin level. Then the incubation media was changed to Krebs buffer containing 20mM glucose. Clusters were incubated for 1h and supernatant was collected for detecting insulin levels. An additional wash was performed between 20mM and 2.8mM to remove residual glucose, followed by repeating this sequence one more time. Finally, a depolarization challenge was performed by incubating the clusters in Krebs buffer containing 2.8 mM glucose and 30 mM KCl for 1 h, and then the supernatant was collected. Clusters were then dissociated using TrypLE Express (Life Technologies) and cells were counted by an automated Vi-Cell (Beckman Coulter). All incubations were carried out at 37°C, with supernatant samples collected at the end of each incubation.

Insulin levels in serum/supernatant samples containing secreted insulin were measured by a Human Ultrasensitive Insulin ELISA kit (ALPCO Diagnostics; 80-INSHUU-E10) followed the manufacturer’s protocol.

### Immunohistochemistry

The testes/kidney of SC-islets transplanted mice were excised and fixed in 4% paraformaldehyde (PFA) in PBS at 4°C overnight. Paraffin embedding and section cutting were done by the HSCI histology core. Before staining, paraffin-embedded slides were treated with Histo-Clear to remove the paraffin. All slides were rehydrated via an ethanol gradient, and the antigen was fixed by incubating in boiling antigen retrieval reagent (10 mM sodium citrate, pH 6.0) for 50 min. For staining, slides were incubated in 5% donkey serum blocking media for 1h, followed with primary antibodies listed above overnight at 4 °C, washed three times, incubated in secondary antibody for 2 h at room temperature, washed, mounted in Vectashield with DAPI (Vector Laboratories; H-1200), covered with coverslips and sealed with clear nail polish. Representative regions were imaged using Zeiss LSM 880 microscopes. The images shown are representative of similar results in at least three biologically separate differentiations from matched or similar stages.

### Mouse Sertoli cell isolation, spleen mononuclear cell isolation, and co-culture experiments

Mouse Sertoli cells need to be isolated and seeded into the gelatin-coated plates one day before setting up for co-culturing. The Sertoli cells were isolated as previously described(*37*). Briefly, the testis was removed and washed twice with PBS. The seminiferous tubules were disseminated in collagenase IV (Gibco, 17104019, 1mg/ml in PBS) at 37°C for 10-15 min. Let the seminiferous tubules settle down and discard the supernatant containing interstitial cells. The seminiferous tubules were then washed 3 times with PBS and moved into 0.25% trypsin solution, incubated at 37°C for 5-10 min, pipette gently until the tubules got fragmented. The single cells were quenched with DMEM with 10%FBS. Sertoli cells and germ cells are pelleted through centrifuge and seeded on the gelatin-coated plate at 0.2-0.3 million/cm^2^, incubated at 37°C overnight. The next day, the Sertoli cells were attached to the bottom of the plates, then remove the supernatant containing germ cells.

Mouse primary spleen mononuclear cells (MNCs) were isolated using C57BL/6 mice. Red blood cells are removed by RBC Lysis Buffer (Biolegend # 420301). MNCs were counted and placed on ice until further use.

### T cell activation assay

Single SC-islet cells are used as target cells. The immune/target cell ratio for co-culture is 1:1. After a 48-hour co-culture, CD4^+^ and CD8^+^ T cells were stained for naïve T cell marker CD62L or T cell activation markers CD69 and CD25.

### T cell proliferation assay

The MNCs were prelabeled with CellTrace™ CFSE dye (Life Technology, C34554) as instructed. After labeling, the immune/target cells were co-cultured in a ratio of 1:1. 3 or 5 days after co-culture. The CD4^+^ cells were analyzed through the 488 channel.

### T cell stimulation(expansion) assay

The CD3/CD28 Dynabeads (Gibco, 11452D) were used to stimulate T cell activation and proliferation. In the stimulation condition, CD4^+^/CD8^+^ cells or cytokines were added to test their immunosuppressive function.

### The Treg-Specific Demethylated Region (TSDR) analysis of CD4^+^/CD8^+^ cells

CD4^+^/CD8^+^ cells and CD4^+^/CD8^-^ cells were isolated from the testis through flow cytometry at 2, 4, 8 weeks after testis transplantation. Treg control was isolated from mouse spleen by Dynabeads™ FlowComp™ Mouse CD4^+^CD25^+^ Treg Cells Kit (life Technology, 11463D). All the cell samples were sent to EpigeneDx to do TSDR analysis.

### The bulk RNA seq of the T_regs_, CD4^+^/CD8^+^ cells, and CD4^+^/CD25^-^ T cells

The T_regs_ and the CD4^+^/CD25^-^ T cells were isolated from C57BL/6 mouse spleen using the Mouse CD4^+^CD25^+^ Treg Cells Kit (Invitrogen 11463D). The CD4^+^/CD8^+^ cells were sorted from testis, which received SC-islets. Each cell type had two biological replicates. All the samples were sent to Bauer Core Facility at Harvard University for further processing and sequencing.

Bulk RNA sequencing data was obtained from two biological duplicates of each cell type and counts of reads mapping to genomic features were produced using the RSEM package. Reads were aligned to mouse genome GRCm39 using the rsem-calculate-expression module with the bowtie2 option and collected using rsem-generate-data-matrix. Expression data were log2 normalized and reference profiles of Treg and CD4 T-cells were obtained by taking the means of the biological replicates. To compare the expression profiles across samples, the 50 most enriched markers were selected from each of the two reference profiles and visualized using a heatmap.

### Cytokine array of the testis supernatant

The digestion of mouse testis cells was described above. 48 hours after culturing, the testis supernatant and the blank medium were collected for the cytokine array (R&D System, ARY006).

### Kidney Capsule transplantation of the SC-islets

Kidney Capsule transplantation of the SC-islets was carried out as previously described^1^ with minor modification. For the CD4^+^/CD8^+^ cells co-transplantation assay, SC-islet clusters were reaggregated with CD4^+^/CD8^+^ cells or CD4^+^/CD8^-^ cells at 10:1 ratio. About 3 x 10^6^ reaggregated clusters were transplanted into male C57BL/6 mice kidney capsules (Charles River). For the cytokine co-transplantation assay, about 5 × 10^6^ SC-islet clusters were embedded into Matrigel. We used a low dose of IL-2 as the same as the most efficient dose to induce T_regs_ when used in the T1D treatment clinical trial (0.47×10^6^IU/m^2^)(*38*) (R&D, 402-ML-020/CF), 100 μg/ml mouse recombinant IL-10(R&D, 417-ML-025/CF), and 100 μg/ml mouse recombinant TGFβ1(R&D, 7666-MB-005/CF) were also added into Matrigel and transplanted together into the kidney capsule of male C57BL/6 mice (Charles River) aged between 8 and 12 weeks. At the specified time after transplantation, kidneys containing grafts were dissected and fixed in 4% PFA overnight at 4 °C. The fixed kidneys were embedded in paraffin and sectioned for immunofluorescence staining, which was performed as described above. All animal studies were approved by the Harvard University IACUC.

### *In vivo* AAV transduction

ssAAV plasmids encoding Firefly luciferase (Luc2) (control) and Luc2-IL2N88D-TGFβ (experimental) (Fig. 6A) were packaged in AAV8-Y447F+Y773F(*39*) (PackGene) and injected i.p. into 4-week-old female NOD mice (n=10/group) (Jackson Laboratories) at a concentration of 5×1011 GC/mouse. Blood glucose and body weight measurements were recorded weekly until blood glucose measured >200 mg/dL, after which measurements were recorded daily. *In vivo* bioluminescence imaging was performed weekly after i.p. administration of 10 μL/g D-luciferin (15 mg/ml) (GoldBio) and imaged 10 min post-injection on the IVIS Spectrum (PerkinElmer).

## Acknowledgments

We would like to thank Stephan Kissler and Ramona Pop for the discussions on manuscript content.

## Funding

Harvard Stem Cell Institute DP-0180-18-04 (DAM)

Juvenile Diabetes Research Foundation 5-COE-2020-967-M-N (DAM)

JPB Foundation (Award #2695)(DAM)

Crown Prince Court Abu Dhabi, UAE and the Howard Hughes Medical Institute (DAM)

## Author contribution

QZ conceived the study.

The experiments were performed by QZ and HL.

IN provided technical support and conceptual advice.

JZ, MH, and ER produced SC-islets for the experiments.

DG, XW, and JKR were involved in the experimental design and advised on the project.

QZ and DAM wrote the manuscript.

DAM designed and supervised the research.

---

**Figure S1.**
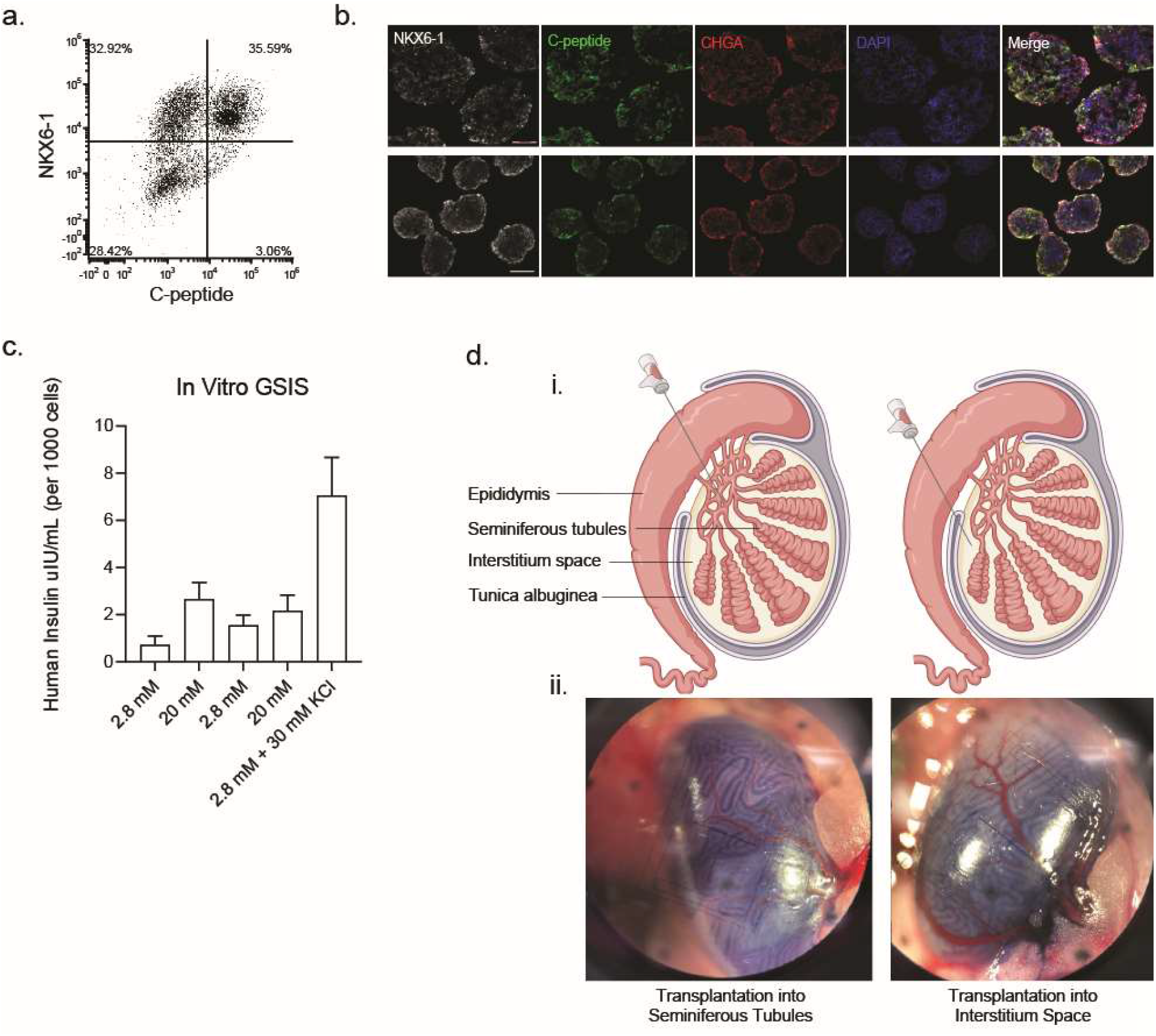
*In vitro* differentiation of SC-islets from human ES cells. A. Flow cytometry shows NKX6-1 and C-peptide expression for beta cells in Stage 6 of differentiation. B. Immunostaining of NKX6-1, C-peptide, and Chromogranin A before and after cluster resizing to 100μm. C. *In vitro* GSIS of SC-islets shows function. D. Transplantation site in the testis, in the seminiferous tubules (left), in the interstitium space of the testis(right)

**Figure S2.**
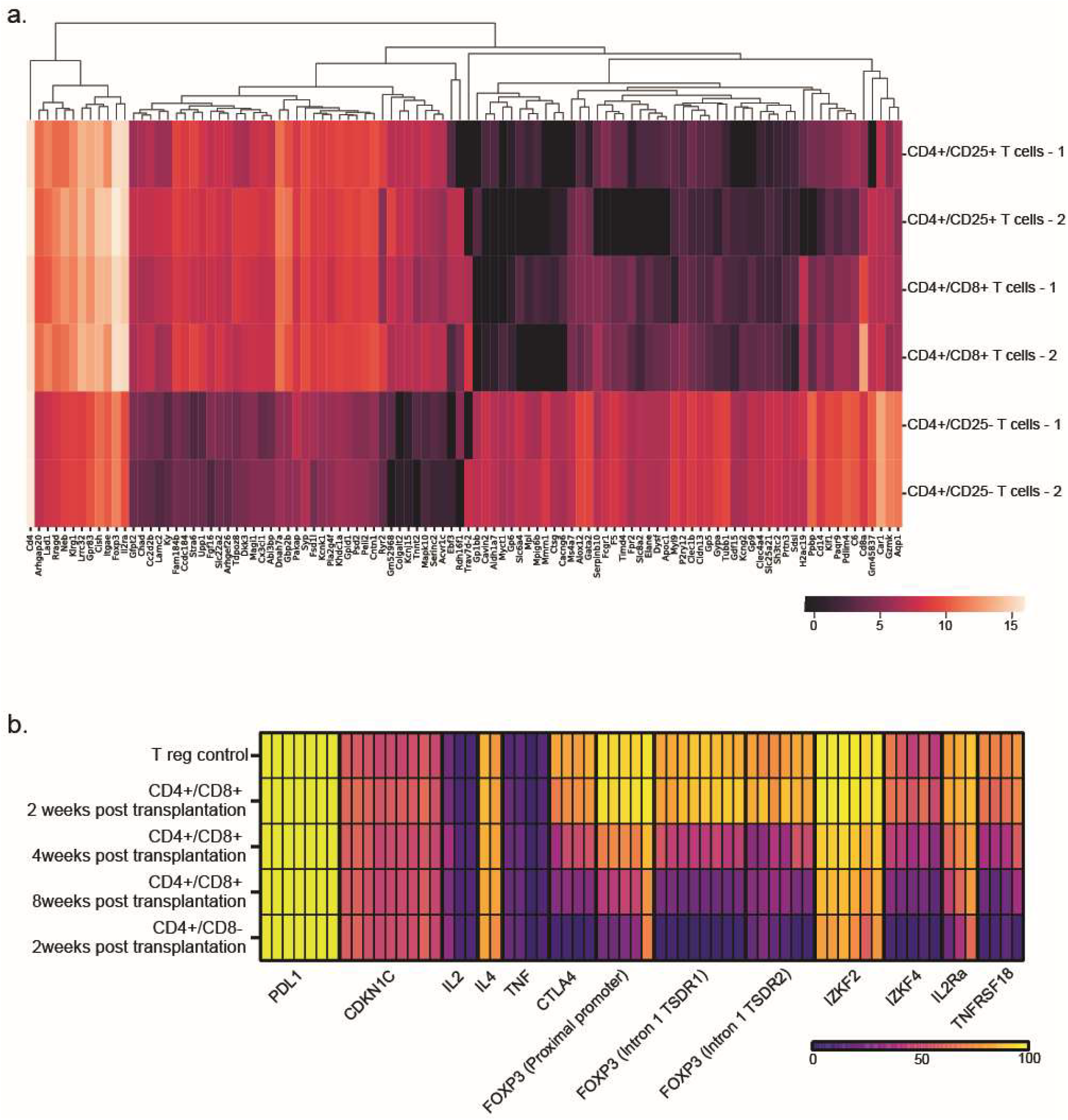
Identification of CD4^+^/CD8^+^ population in testis transplantation A. Bulk RNA seq of CD4^+^/CD25^+^ T_regs_, CD4^+^/CD8^+^ cell population, and CD4^+^/CD25^-^ cell population. The top 100 differentially expressed genes in CD4^+^/CD25^+^ T_regs_ and CD4^+^/CD25^-^ cell population are shown B. Bisulfite sequencing of gene methylation levels of T cell-related genes for T_regs_ control, CD4^+^/CD8^-^ cells, and CD4^+^/CD8^+^ cells sorted from testis at 2, 4, and 8 weeks after transplantation.

## References

1. F. W. Pagliuca et al., Generation of functional human pancreatic beta cells in vitro. Cell 159, 428–439 (2014).

2. Q. P. Peterson et al., A method for the generation of human stem cell-derived alpha cells. Nat Commun 11, 2241 (2020).

3. A. Veres et al., Charting cellular identity during human in vitro beta-cell differentiation. Nature 569, 368–373 (2019).

4. A. Rezania et al., Production of functional glucagon-secreting alpha-cells from human embryonic stem cells. Diabetes 60, 239–247 (2011).

5. H. A. Russ et al., Controlled induction of human pancreatic progenitors produces functional beta-like cells in vitro. EMBO J 34, 1759–1772 (2015).

6. X. Han et al., Generation of hypoimmunogenic human pluripotent stem cells. Proc Natl Acad Sci U S A 116, 10441–10446 (2019).

7. H. Xu et al., Targeted Disruption of HLA Genes via CRISPR-Cas9 Generates iPSCs with Enhanced Immune Compatibility. Cell Stem Cell 24, 566–578 e567 (2019).

8. T. Deuse et al., Hypoimmunogenic derivatives of induced pluripotent stem cells evade immune rejection in fully immunocompetent allogeneic recipients. Nat Biotechnol 37, 252–258 (2019).

9. C. J. Taylor, S. Peacock, A. N. Chaudhry, J. A. Bradley, E. M. Bolton, Generating an iPSC bank for HLA-matched tissue transplantation based on known donor and recipient HLA types. Cell Stem Cell 11, 147–152 (2012).

10. S. Zhao, W. Zhu, S. Xue, D. Han, Testicular defense systems: immune privilege and innate immunity. Cell Mol Immunol 11, 428–437 (2014).

11. N. Li, T. Wang, D. Han, Structural, cellular and molecular aspects of immune privilege in the testis. Front Immunol 3, 152 (2012).

12. C. Y. Cheng, D. D. Mruk, The blood-testis barrier and its implications for male contraception. Pharmacol Rev 64, 16–64 (2012).

13. J. R. Head, W. B. Neaves, R. E. Billingham, Immune privilege in the testis. I. Basic parameters of allograft survival. Transplantation 36, 423–431 (1983).

14. M. Huo et al., Role of IL-17 Pathways in Immune Privilege: A RNA Deep Sequencing Analysis of the Mice Testis Exposure to Fluoride. Sci Rep 6, 32173 (2016).

15. G. Kaur, S. Vadala, J. M. Dufour, An overview of a Sertoli cell transplantation model to study testis morphogenesis and the role of the Sertoli cells in immune privilege. Environ Epigenet 3, dvx012 (2017).

16. O. Boyman, M. Kovar, M. P. Rubinstein, C. D. Surh, J. Sprent, Selective stimulation of T cell subsets with antibody-cytokine immune complexes. Science 311, 1924–1927 (2006).

17. M. A. Caligiuri et al., Selective modulation of human natural killer cells in vivo after prolonged infusion of low dose recombinant interleukin 2. J Clin Invest 91, 123–132 (1993).

18. D. A. Horwitz, S. G. Zheng, J. Wang, J. D. Gray, Critical role of IL-2 and TGF-beta in generation, function and stabilization of Foxp3+CD4+ Treg. Eur J Immunol 38, 912–915 (2008).

19. K. W. Moore, R. de Waal Malefyt, R. L. Coffman, A. O’Garra, Interleukin-10 and the interleukin-10 receptor. Annu Rev Immunol 19, 683–765 (2001).

20. A. O’Garra, P. Vieira, Regulatory T cells and mechanisms of immune system control. Nat Med 10, 801–805 (2004).

21. L. B. Peterson et al., A long-lived IL-2 mutein that selectively activates and expands regulatory T cells as a therapy for autoimmune disease. J Autoimmun 95, 1–14 (2018).

22. L. Khoryati et al., An IL-2 mutein engineered to promote expansion of regulatory T cells arrests ongoing autoimmunity in mice. Sci Immunol 5, (2020).

23. M. Bagot et al., Isolation of tumor-specific cytotoxic CD4+ and CD4+CD8dim+ T-cell clones infiltrating a cutaneous T-cell lymphoma. Blood 91, 4331–4341 (1998).

24. J. Desfrancois et al., Double positive CD4CD8 alphabeta T cells: a new tumor-reactive population in human melanomas. PLoS One 5, e8437 (2010).

25. G. Sarrabayrouse et al., Tumor-reactive CD4+ CD8alphabeta+ CD103+ alphabetaT cells: a prevalent tumor-reactive T-cell subset in metastatic colorectal cancers. Int J Cancer 128, 2923–2932 (2011).

26. Y. Parel et al., Presence of CD4+CD8+ double-positive T cells with very high interleukin-4 production potential in lesional skin of patients with systemic sclerosis. Arthritis Rheum 56, 3459–3467 (2007).

27. G. Das et al., An important regulatory role for CD4+CD8 alpha alpha T cells in the intestinal epithelial layer in the prevention of inflammatory bowel disease. Proc Natl Acad Sci U S A 100, 5324–5329 (2003).

28. M. Szczepanik et al., Epicutaneous immunization induces alphabeta T-cell receptor CD4 CD8 double-positive non-specific suppressor T cells that inhibit contact sensitivity via transforming growth factor-beta. Immunology 115, 42–54 (2005).

29. Q. Z. Dario Gerace, Jennifer Hyoje-Ryu Kenty, Elad Sintov, Xi Wang, Kyle R Boulanger, Hongfei Li, Douglas A Melton, Secreted cytokines provide local immune tolerance for human stem cell-derived islets. Biorxiv, (2022).

30. J. Koreth et al., Interleukin-2 and regulatory T cells in graft-versus-host disease. N Engl J Med 365, 2055–2066 (2011).

31. D. Saadoun et al., Regulatory T-cell responses to low-dose interleukin-2 in HCV-induced vasculitis. N Engl J Med 365, 2067–2077 (2011).

32. A. Hartemann et al., Low-dose interleukin 2 in patients with type 1 diabetes: a phase 1/2 randomised, double-blind, placebo-controlled trial. Lancet Diabetes Endocrinol 1, 295–305 (2013).

33. M. Rosenzwajg et al., Low-dose interleukin-2 fosters a dose-dependent regulatory T cell tuned milieu in T1D patients. J Autoimmun 58, 48–58 (2015).

34. J. A. Todd et al., Regulatory T Cell Responses in Participants with Type 1 Diabetes after a Single Dose of Interleukin-2: A Non-Randomised, Open Label, Adaptive Dose-Finding Trial. PLoS Med 13, e1002139 (2016).

35. Y. Yuan et al., Generation of fertile offspring from Kit(w)/Kit(wv) mice through differentiation of gene corrected nuclear transfer embryonic stem cells. Cell Res 25, 851–863 (2015).

36. S. Chuma et al., Spermatogenesis from epiblast and primordial germ cells following transplantation into postnatal mouse testis. Development 132, 117–122 (2005).

37. M. D. Anway, J. Folmer, W. W. Wright, B. R. Zirkin, Isolation of sertoli cells from adult rat testes: an approach to ex vivo studies of Sertoli cell function. Biol Reprod 68, 996–1002 (2003).

38. E. Seelig et al., The DILfrequency study is an adaptive trial to identify optimal IL-2 dosing in patients with type 1 diabetes. JCI Insight 3, (2018).

39. M. Chen et al., Efficient Gene Delivery and Expression in Pancreas and Pancreatic Tumors by Capsid-Optimized AAV8 Vectors. Hum Gene Ther Methods 28, 49–59 (2017).

